# Frontal cortex tracks surprise separately for different sensory modalities but engages a common inhibitory control mechanism

**DOI:** 10.1101/572081

**Authors:** Jan R. Wessel, David E. Huber

## Abstract

The brain constantly generates predictions about the environment to guide action. Unexpected events lead to surprise and can necessitate the modification of ongoing behavior. Surprise can occur for any sensory domain, but it is not clear how these separate surprise signals are integrated to affect motor output. By applying a trial-to-trial Bayesian surprise model to human electroencephalography data recorded during a cross-modal oddball task, we tested whether there are separate predictive models for different sensory modalities (visual, auditory), or whether expectations are integrated across modalities such that surprise in one modality decreases surprise for a subsequent unexpected event in the other modality. We found that while surprise was represented in a common frontal signature across sensory modalities (the fronto-central P3 event-related potential), the single-trial amplitudes of this signature more closely conformed to a model with separate surprise terms for each sensory domain. We then investigated whether surprise-related fronto-central P3 activity indexes the rapid inhibitory control of ongoing behavior after surprise, as suggested by recent theories. Confirming this prediction, the fronto-central P3 amplitude after both auditory and visual unexpected events was highly correlated with the fronto-central P3 found after stop-signals (measured in a separate stop-signal task). Moreover, surprise-related and stopping-related activity loaded onto the same component in a cross-task independent components analysis. Together, these findings suggest that medial frontal cortex maintains separate predictive models for different sensory domains, but engages a common mechanism for inhibitory control of behavior regardless of the source of surprise.

**Author summary:** Surprise is an elementary cognitive computation that the brain performs to guide behavior. We investigated how the brain tracks surprise across different senses: Do unexpected sounds make subsequent unexpected visual stimuli less surprising? Or does the brain maintain separate expectations of environmental regularities for different senses? We found that the latter is the case. However, even though surprise was separately tracked for auditory and visual events, it elicited a common signature over frontal cortex in both sensory domains. Importantly, we observed the same neural signature when actions had to be stopped after non-surprising stop-signals in a motor inhibition task. This suggests that this signature reflects a rapid interruption of ongoing behavior when our surroundings do not conform to our expectations.

## 1. Introduction

Surprise occurs when expectations about the multi-sensory environment are violated. It provides an elementary cognitive and physiological process that forms the backbone of many influential theories of cognitive processing and control (1-5). The rapid modification of ongoing actions after surprise is critical for effective goal-directed behaviors (6, 7). For example, while eating berries, one needs to rapidly stop ongoing actions when encountering a berry that looks, smells, or feels surprising, lest one eats a rotten berry. However, the manner in which the brain tracks surprise across different sensory domains is not fully understood.

Prior imaging work has shown that unexpected events, regardless of their sensory modality, activate similar brain networks (8-11). In line with this, scalp-electroencephalography (EEG) shows that unexpected events are followed by a modality-independent fronto-central P3 event-related potential (12, ERP, 13). The canonical neural response to surprise across modalities could indicate that the brain integrates environmental information across modalities and generates global predictions that form the basis of surprise-processing. Alternatively, surprise might result from separate, independent predictions for each sensory domain. In this latter case, the modality-independent surprise response could index a common set of downstream mechanisms triggered by surprise, regardless of sensory domain.

In the current study, we tested these two alternatives against each other. While performing a cross-modal oddball task (CMO, 14), human subjects were presented with visual or auditory unexpected events. Using the statistics of the trial sequence, we constructed two models of Bayesian surprise (5). In one model, surprise-values were separately coded for each sensory domain (i.e., an unexpected sound did not reduce surprise of a subsequent unexpected visual event). In the alternative model, surprise was coded in a common term across modalities (i.e., an unexpected sound reduced surprise for a subsequent unexpected visual event). We fit both models to the trial-to-trial electroencephalographic response to unexpected events at each of 64 scalp-sites to determine which model better represents the neural surprise response.

As mentioned above, in case this trial-to-trial modeling of the neural surprise response suggests that surprise-terms are computed separately for each sensory domain (i.e., surprise is not integrated into a common model), the expected cross-modal overlap in neural response may be explained by a common, supra-modal control mechanism that is triggered by surprise, regardless of modality. Therefore, in a second step, we aimed to test the hypothesis that the fronto-central P3 after unexpected events indexes the modality-independent activation of a cognitive control mechanism aimed at inhibiting ongoing behavior. This hypothesis was recently proposed in a theoretical framework claiming that surprise automatically engages the same motor inhibition mechanism that is recruited when ongoing actions have to be stopped (15). The activity of this mechanism can be measured in the stop-signal task (SST, 16), where fronto-central P3 activity following (non-surprising) stop-signals indexes the speed of motor inhibition (17, 18). To determine whether the fronto-central P3 after unexpected events in the CMO task and the P3 after stop-signals in the SST reflect the same process, we first correlated their amplitudes across tasks and subjects. We hypothesized that if they indeed reflect the same process, their amplitudes should be positively correlated. Additionally, we used independent component analysis to determine if both fronto-central waveforms load onto a common independent component (19, 20). In doing so, we aimed to provide converging support for the proposal that surprise-signals in frontal cortex lead to the automatic activation of a control process that aims to inhibit ongoing behavior, independent of the modality of the unexpected event.

## 2. Materials and Methods

### 2.1. Participants

Fifty-five healthy young adult volunteers from the Iowa City community were recruited via a research-dedicated email list, as well as through the University of Iowa Department of Psychological and Brain Sciences’ online subject recruitment tool. The sample consisted of thirty-one females and twenty-four males (mean age: 20.9 y, SEM: 0.05, range 18-31), eight of them left-handed. Participants were compensated with course credit or an hourly payment of $15. The procedure was approved by the University of Iowa Institutional Review Board (#201612707).

### 2.2. Materials

Stimuli for both tasks were presented using the Psychophysics toolbox (21) (RRID:SCR_002881) under MATLAB 2015b (TheMathWorks, Natick, MA; RRID:SCR_001622) on an IBM-compatible computer running Fedora Linux. Visual stimuli were presented on an ASUS VG278Q low-latency flat screen monitor, while sounds were played at conversational volume through speakers positioned on either side of the monitor. Responses were made using a standard QWERTY USB-keyboard.

### 2.3. Cross-Modal Oddball task

Each trial began with a central white fixation cross on black background (500ms), which was followed by an audio-visual cue (Figure 1). Participants were instructed that this cue would be informative regarding the timing of a subsequent target stimulus (a left-or rightward white arrow) that they would have to respond to. Participants were instructed that the cue would consist of a green circle presented in place of the fixation cross for 200ms, accompanied by a 600Hz sine wave tone of 200ms duration. After cue presentation, the fixation cross reappeared for 300ms, followed by the target (i.e., the target appeared exactly 500ms after cue onset). Participants were instructed to respond to the target as fast as possible. Target responses were collected through the keyboard (q for leftward and p for rightward arrows) with the index finger of the respective hand. Participants had 1,000ms to respond to the target, after which the fixation cross reappeared and the inter-trial interval began. The duration of the inter-trial interval lasted until 2,500ms from the initial onset of the fixation cross (beginning of the trial) was reached. Furthermore, to prevent predictable trial initiation timing, a variable-length jitter was added to the ITI (100 – 500ms in 100ms increments, uniform distribution), resulting in an overall trial duration ranging from 2,600ms to 3,000ms.

**Figure 1.**
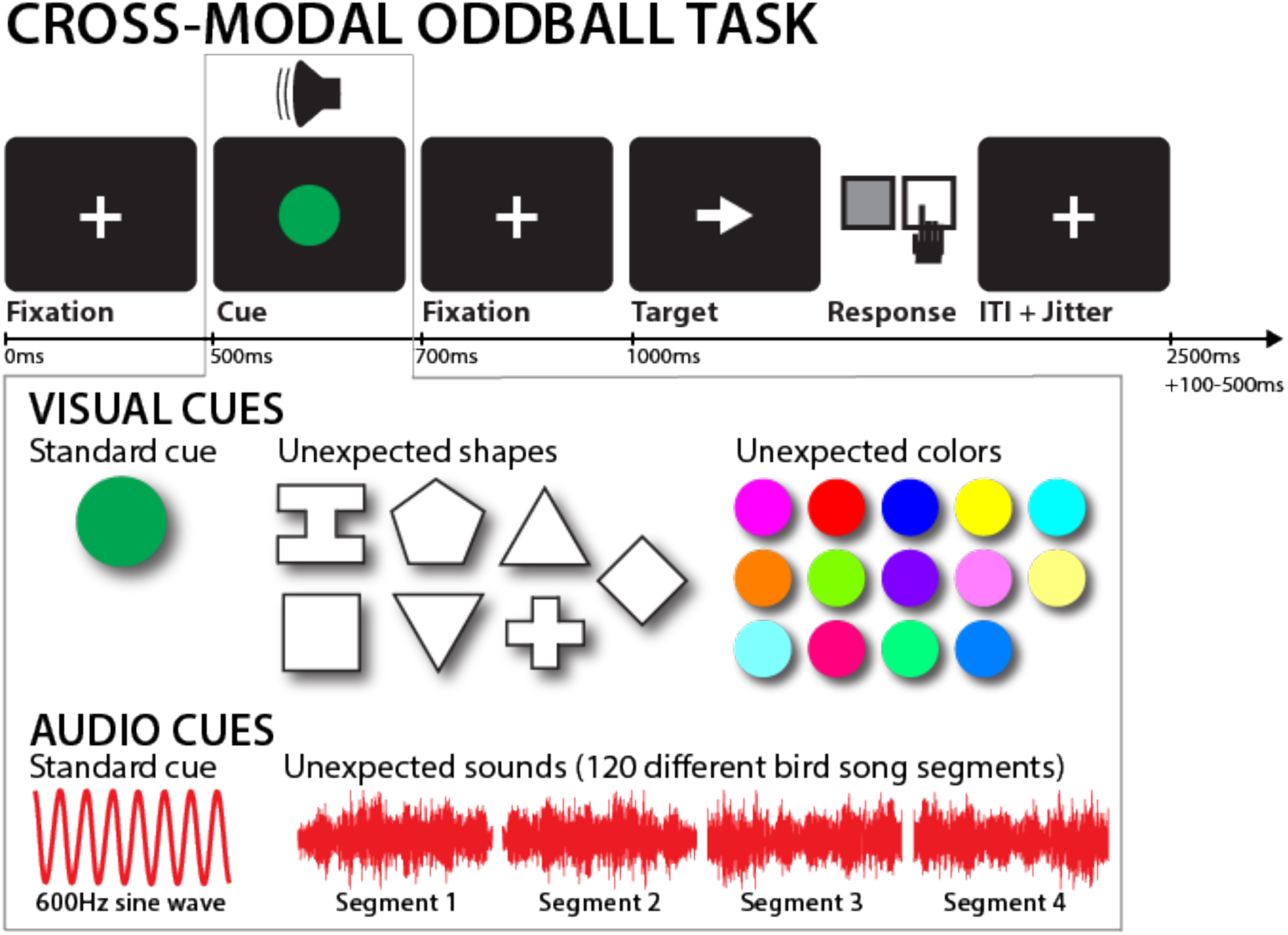
Cross-modal oddball task diagram. The top row depicts the trial timing. The gray box attached to the cue illustrates the different cue properties by trial type. Each cue consisted of a visual and an auditory component. Standard visual cues consisted of a green circle, whereas unexpected visual cues were one of seven non-circular shapes shown in one of fourteen non-green colors. Standard auditory cues consisted of a 600Hz sine wave, whereas unexpected auditory cues were one of 120 individual unique birdsong segments. On a trial that contained an unexpected cue in one domain, the part of the cue always contained the standard component.

After 10 practice trials without any unexpected cues, participants performed 240 trials, spread across 4 blocks. During this experimental trials, 80% of trials contained cues that were as described above (hereafter referred to as standard cues). On 10% of trials, the sine-wave tone was replaced with one of 120 unique birdsong segments, which were matched in amplitude and duration to the sine-wave tone (unexpected auditory cue). For these auditory unexpected cues, the visual part of the cue remained the same as for standard trials. On the remaining 10% of trials, the green circle was replaced by one of seven different geometric shapes (upwards/downwards triangle, square, diamond, cross, hexagon, or a serifed “I”-shape) in one of 15 different non-green colors spread across the RGB spectrum (unexpected visual cue, cf. Figure 1). For these visual unexpected cues, the auditory part of the cue remained the same as for standard trials. Trials were presented in pseudorandom order, with the following constraints: the three first trials of each block had to contain standard cues; no two consecutive unexpected-cue trials were allowed to occur; and each block had to have the same number of unexpected auditory and visual cues.

### 2.4. Stop-signal task

Trials began with a white fixation cross on a gray background (500ms duration), followed by a white leftward- or rightward-pointing arrow (go-signal). Participants had to respond as fast and accurately as possible to the arrow by using their left or right index finger as indicated by the direction of the arrow (the respective response-buttons were q and p on the QWERTY keyboard). On 33% of trials, a stop-signal occurred (the arrow turned from white to red) at a delay after the go-stimulus (stop-signal delay, SSD). The SSD, which was initially set to 200ms, was dynamically adjusted in 50ms increments to achieve a p(stop) of .5: after successful stops, the SSD was increased; after failed stops, it was decreased. This was done independently for leftward and rightward go-stimuli: SSD started at 200ms for both left- and right-arrow trials. Then, if a stop-trial with a leftward arrow lead to a failed stop, the SSD for the next leftward arrow was decreased by 50ms, whereas the SSD for the next rightward response remained unchanged. This way, the SSD was allowed to vary independently for each arrow/response direction. Trial duration was fixed at 3000ms. Six blocks of 50 trials were performed (200 go, 100 stop). Before the main experiment, subjects practiced the task for 24 trials (16 go, 8 stop).

### 2.5. Code availability

All analysis code, as well as the task code, can be downloaded alongside the raw data at the following URL: [to be inserted upon acceptance].

### 2.6. Behavioral analysis

For the CMO task, we quantified mean reaction time (RT), mean error rate (wrong button pressed), and mean miss rate (no response made within 1,000ms after target onset) for each of the three trial types (standard cue, unexpected auditory cue, unexpected visual cue). We analyzed these dependent variables using a 3 x 4 repeated-measures ANOVA with the factors TRIAL TYPE (1-3) and BLOCK (1-4). In case of a significant interaction, we performed follow-up paired-samples t-tests that compared each of the two unexpected cue conditions to the standard cue condition separately for each of the four blocks, resulting in eight total tests. The alpha-level for these comparisons was corrected using the Bonferroni correction to a corrected alpha of .0063 (i.e., p = .05 / 8).

For the stop-signal task, we examined the following measures: mean Go-trial RT, mean failed-stop trial RT, and mean stop-signal RT (SSRT; computed using the integration method, Verbruggen & Logan, 2009; Boehler et al., 2014).

### 2.7. EEG recording

EEG was recorded using a 62-channel electrode cap connected to two BrainVision MRplus amplifiers (BrainProducts, Garching, Germany). Two additional electrodes were placed on the left canthus (over the lateral part of the orbital bone of the left eye) and over the part of the orbital bone directly below the left eye. The ground was placed at electrode Fz, and the reference was placed at electrode Pz. EEG was digitized at a sampling rate of 500 Hz.

### 2.8. EEG preprocessing

The CMO and SST datasets were preprocessed separately, using custom routines in MATLAB, incorporating functions from the EEGLAB toolbox (22). The channel * time-series matrices for each task were imported into MATLAB and then filtered using symmetric two-way least-squares finite impulse response filters (high-pass cutoff: .3 Hz, low-pass cutoff: 30 Hz). Non-stereotyped artifacts were automatically removed from further analysis using segment statistics applied to consecutive one-second segments of data (joint probability and joint kurtosis, with both cutoffs set to 5 SD, cf., 23). After removal of non-stereotypic artifacts, the data were then re-referenced to common average and subjected to a temporal infomax ICA decomposition algorithm (24), with extension to subgaussian sources (25). The resulting component matrix was screened for components representing eye-movement and electrode artifacts using outlier statistics and non-dipolar components (residual variance cutoff at 15%, 26), which were removed from the data. The remaining components (an average of 17.1 per subject) were subjected to further analyses.

### 2.9. Experimental design and statistical tests (EEG analysis)

#### 2.9.1. Hypothesis 1 – Cross-modal representation of surprise

To investigate whether surprise is represented separately for each sensory domain, we constructed two Bayesian surprise terms on a trial-by-trial basis, based on the trial sequences for each subject (cf. Figure 2). For both terms, the surprise value associated with an unexpected cue on a particular trial was based on the following equation:

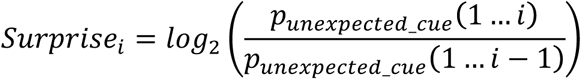

**Figure 2.**
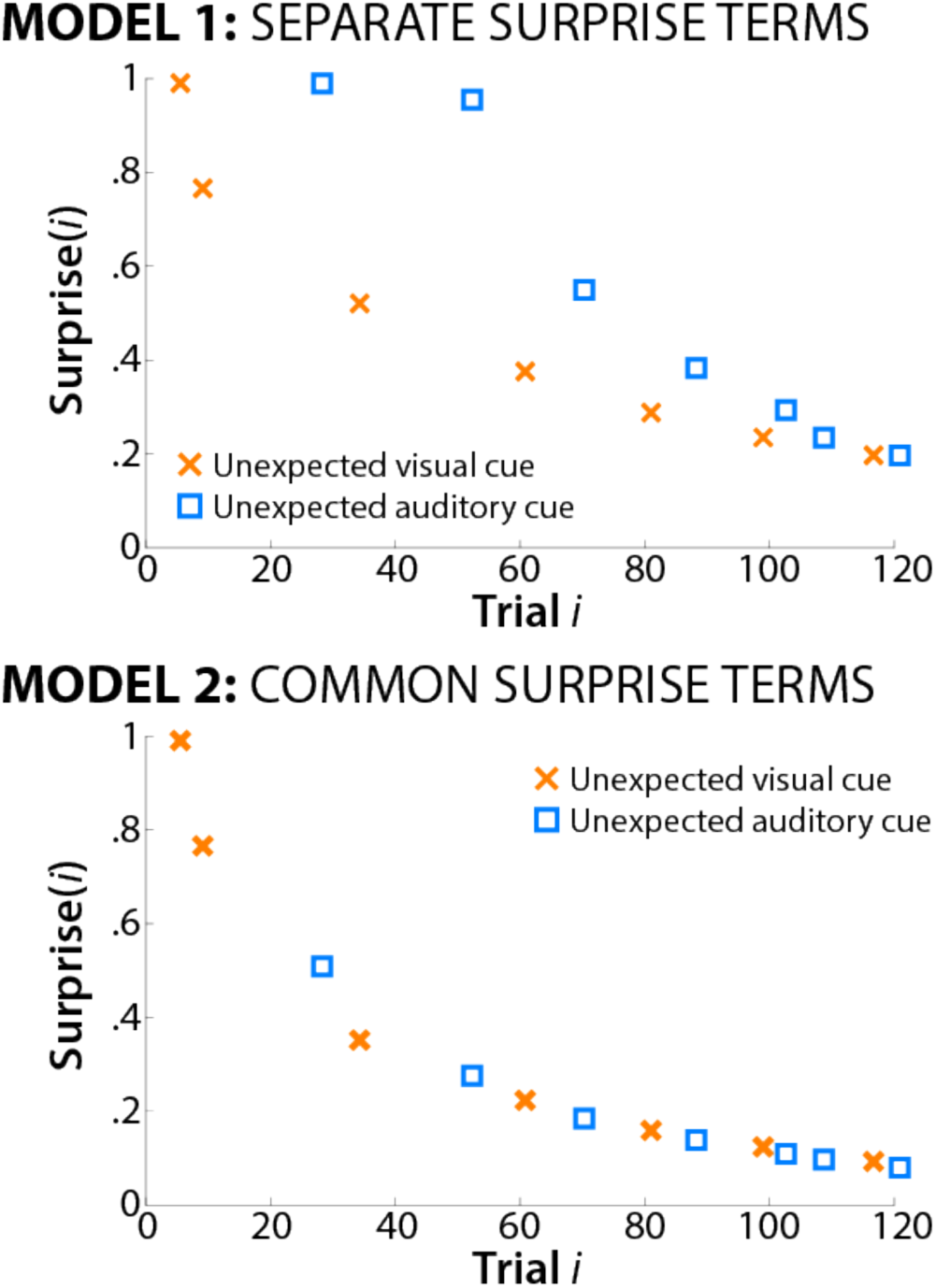
Single-subject example of surprise-term construction for each model. Top: Model 1 uses separate surprise terms for each sensory domain. In effect, the presence of a surprising event in one sensory domain does not inform the prior in the other sensory domain. Bottom: Model 2 uses a combined surprise term across both domains. In effect, all unexpected events, regardless of domain, influence the construction of the prior.

This equation corresponds to the trial-wise Kullback-Leibler divergence between the prior probability of an unexpected cue (denominator) and the posterior probability of an unexpected cue (numerator). This value is bounded between 0 (posterior = prior -> no surprise) and 1 (maximum surprise). Since this value is not defined on the first occurrence of an unexpected cue (where the prior is zero, leading to a division by 0), the surprise value for that trial was set to 1 (maximum surprise).

Based on this equation, we generated two different models. In Model 1 (separate surprise terms, Figure 2A), values for each sensory domain were calculated separately. In other words, the first time the subject encountered an unexpected auditory cue in the trial sequence, the surprise for that trial was 1. Subsequent unexpected auditory cues then produced lower surprise values as the posterior and prior probabilities of unexpected auditory cues converge on the same value (i.e., as the ratio approaches 1, the log approaches 0) with increasing numbers of previous unexpected auditory cues. Critically, these prior and posterior probabilities for *auditory* cues are calculated without reference to the number of prior unexpected *visual* cues. Thus, once a subject encounters the first unexpected *visual* cue, the surprise value for that trial is again 1 (maximum surprise). Hence, the prior for each sensory domain is unaffected by the occurrence of unexpected cues in the other sensory domain.^1^

In contrast to Model 1, Model 2 (common surprise term, Figure 2B) extracted a combined surprise value, calculated without reference to sensory domain. In other words, the prior and posterior probabilities are based on the number of unexpected cues, regardless of whether those cues were visual or auditory.

These values were then used to model the whole-brain event-related single-trial EEG response on all trials that contained unexpected cues. This was done using procedures reported by Fischer and Ullsperger (27). For each subject, sixty-four matrices (one for each EEG channel) were generated that contained the event-related EEG response for each individual trial with an unexpected cue (24 auditory, 24 visual = 48), measured in 10 consecutive time windows covering the entire cue-target interval (500ms, Figure 3). The time windows were centered around time points ranging from 50 to 500ms and were 48ms long (24ms before and after the exact time point). EEG activity within each time window was averaged for each trial (prior to averaging, the single-trial data were baseline-corrected by subtracting the activity ranging from 100ms – 0ms relative to the cue). Hence, this resulted in a matrix of 48 (trials) * 10 (time points) for each channel (unless trials were excluded because of artifacts); cf. the blue matrix in Figure 3. Both of the two candidate surprise models constructed from the Bayesian equation were then applied to these EEG matrices. In applying the models, both the surprise terms and EEG response were z-scored (to standardize the resulting beta weights) and the model terms were regressed onto each time-window vector of the trial by time window EEG response matrix. This was done using MATLAB’s robustfit() function, which performs a linear regression that is robust to outliers.

**Figure 3.**
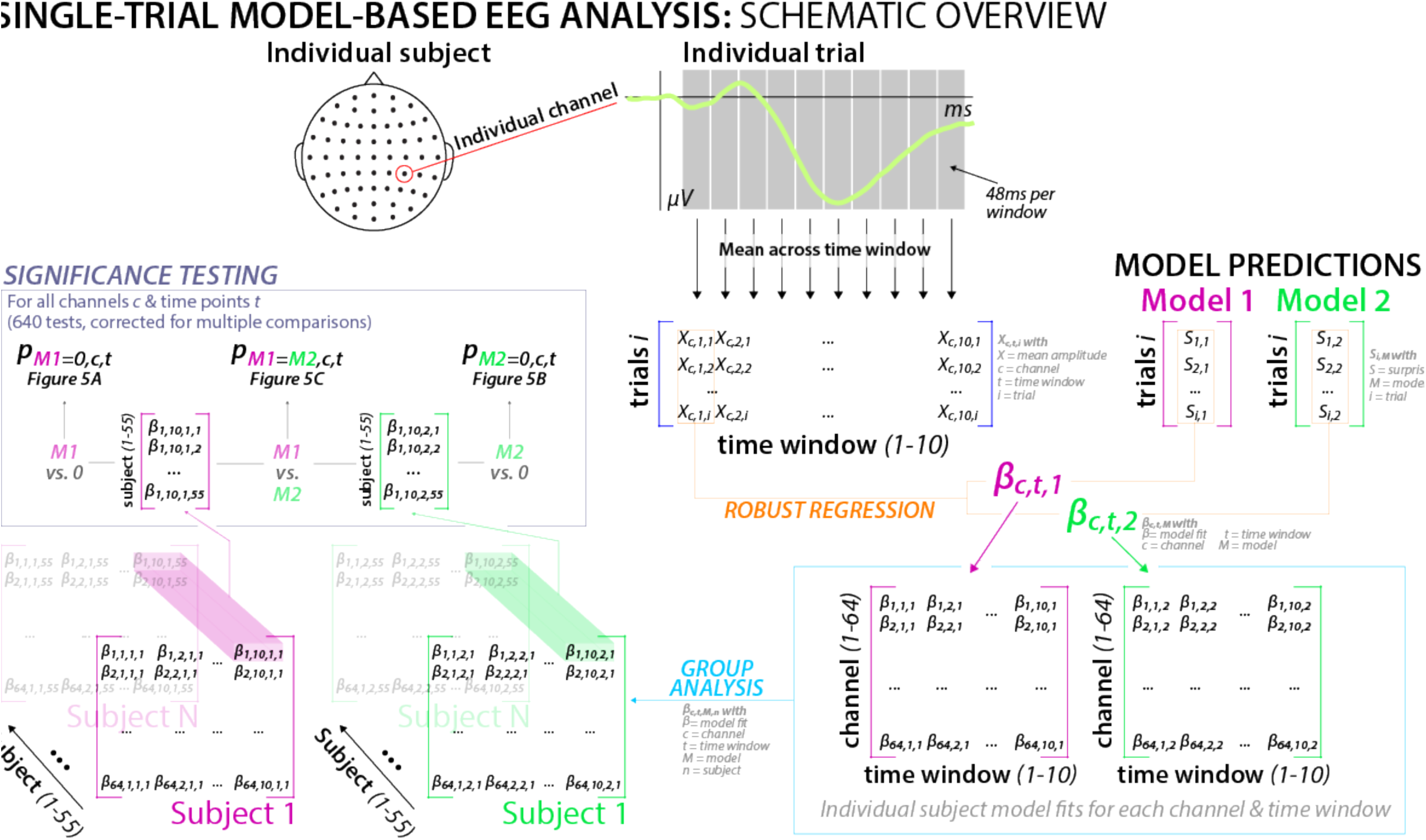
Schematic overview of the single-subject, single-trial robust regression analysis, mapping surprise terms from two models onto the whole brain EEG response to unexpected cues (as well as performing a model comparison). Clockwise from the top-left: For each individual channel, the trial-by-trial event-related response was averaged within 10 consecutive time windows following onset of an unexpected cue. This resulted in a matrix of 48 trials by 10 time windows of EEG amplitude values for each subject (one for each channel; blue brackets). Each subject’s individual model terms for both models (pink and green brackets on the top right) were then correlated with each of the trial-vectors for each time window using robust regression (orange line). The resulting beta values were stored in one channel by time window matrix for each subject and model (bottom right). These beta weights were subjected to group-level analyses across subjects (bottom left), with each channel by time window combination (640 unique combinations per model) tested against 0 for each model separately (purple box), with paired samples t-tests using subject as the random factor.

The resulting matrix of beta values was tested against 0 (using paired-samples t-tests for the beta values, with subject as the random factor) at each channel and time point separately. This identified channels and time periods at which the respective model surprise terms reliably captured variability in the EEG signal. This resulted in two sets of 64 (channels) * 10 (time points) = 640 individual tests (one set for each model). To test which model provided a superior fit of the neural data at each channel and time-point, the resulting beta weights from each model also tested against each other, producing a third set of 640 paired-samples t-test (again with subject as the random factor).

To correct for multiple comparisons across these three sets of 640 t-tests, we adjusted the alpha-level using the false discovery rate correction procedure (FDR, 28) based on a family-wise alpha-level of .01. This resulted in an adjusted alpha-level of p = .00044. A detailed graphical illustration of this overall analysis strategy can be found in Figure 3.

In a separate exploratory analysis of the trial-to-trial reaction times, we similarly regressed the surprise terms from each model onto the response latencies for each target stimulus to assess whether surprise, according to each model, predicted slower responses.

#### 2.9.2. Hypothesis 2 – Surprise-related frontal cortex activity reflects inhibitory control

In addition to our above-described test of whether surprise is represented in the brain separately for each sensory domain, we also tested whether the predicted fronto-central neural response to unexpected cues (i.e., the P3) reflects an inhibitory control signal aimed at inhibiting ongoing behavior during surprise. To this end, we employed cross-task comparisons between the fronto-central P3 extracted for each subject from the CMO task and a separate ‘functional localizer’ task – the stop-signal task – which all subjects performed after the CMO task (subjects performed the SST after the CMO task so they were not biased to use inhibitory control in the CMO task). We used two different approaches to compare activity across tasks: amplitude correlations and indepdendent component analysis (ICA).

##### 2.9.2.1. Amplitude correlations (Approach 1)

In the first approach, we assessed correlations between EEG amplitudes across tasks. More specifically, if the fronto-central signals from each task reflect the same brain process, they should be positively correlated (e.g., a subject with a more pronounced stop-signal P3 should also show a larger P3 to unexpected cues in the CMO task). However, positive correlations might arise from a variety of nuisance variables (e.g., better signal-to-noise ratio for some subjects compared to others), and these alternatives were addressed by comparing these correlations with various control correlations.

To perform our correlation analyses, for each subject, we extracted the amplitudes of several trial-averaged event-related potentials (ERPs) from both tasks, all of which were averaged from -100 to 700ms with respect to the time-locking event (and baseline corrected from -100 to 0ms):

1. ERPs of interest:
  - Fronto-central P3 following the stop-signal on successful stop-trials in the SST
  - Fronto-central P3 following visual or auditory unexpected cues in the CMO task.
2. Control ERPs:
  - posterior occipital (visual) N1 to the arrow stimuli in both tasks (i.e., to the Go-signal in the SST and to the target-arrow in the CMO task).
  - Fronto-central P3 following standard cues in the CMO task.

For all P3 ERPs, amplitudes were extracted by measuring the largest positive deflection in the trial-average during the time-window ranging from 250-500ms following the time-locking event (measured at fronto-central electrodes FCz and Cz). For both N1 ERPs, amplitudes were extracted by measuring the largest negative deflection in the trial average during the time-window ranging from 100-300ms following the time-locking event (measured at occipital electrodes Oz, O1, and O2).

We ran the following correlation analyses using the Pearson correlation coefficient:

###### Main hypothesis

If the fronto-central P3 during surprise and after stop-signals signify the same process, there should be a positive correlation between the stop-signal P3 in the stop-signal task and both the visual and auditory unexpected-cue P3 in the cross-modal oddball task.

###### Control analysis 1

It is widely accepted that the occipital N1 is a visual perception process. Hence, there should be a positive correlation between the posterior-occipital N1 to the go-signal arrow in the SST task and the N1 to the target-arrow stimuli in the CMO task. Both stimuli were visually identical and had the same meaning in both tasks (they instructed a motor response in the according direction of the arrow). This control analysis was run to demonstrate that if two ERPs reflect the same process across tasks, their amplitudes will be correlated.

###### Control analysis 2

The correlation between the stop-signal P3 and the occipital N1 to the go-signa arrow l in the SST was examined to rule out the possibility that subjects show similar amplitudes for ERPs within the same task, even when they reflect different processes.

###### Control analysis 3

The correlation between the stop-signal P3 in the SST and the occipital N1 to the target-arrow in the CMO task was examined to rule out the possibility that subjects show similar amplitudes for different ERPs regardless of task and / or process.

###### Control analysis 4

The correlation between the stop-signal P3 in the SST and the fronto-central P3 to standard cues in the CMO task was examined to rule out the possibility that the stop-signal P3 is positively correlated with the fronto-central P3 to any meaningful task cue, even when that cue is not surprising.

###### Correlation comparison

We predicted that our main hypothesis, as well as our control analysis 1, would yield significant positive correlations. We also predicted that our other control analyses (2-4) would not yield significant correlations. Hence, the latter control analyses involve null hypothesis tests, with unknown statistical power.

Therefore, in addition to performing these control analyses, we directly compared the magnitude of all control correlations against the magnitude of the correlations between the stop-signal P3 and the fronto-central P3s to unexpected cues in the CMO task. This tested the alternative hypotheses that the predicted positive correlation would be significantly larger than the nuisance correlations, thereby providing a direct test of our hypotheses. To do so, we used a bootstrapping approach. First, we inverted the N1 amplitudes so that correlations between any of the six amplitude measures (the four P3s and the two N1s) could be interpreted with the same directionality. There were two correlations that were expected to be significant (the stop-signal P3 versus the CMO P3 and the stop-signal N1 versus the CMO N1) and each of these were compared with the three correlations that were expected to be null (control analyses 2-4 above), resulting in six correlation differences. To test whether these differences were significant, we repeated the same analysis 5000 times, but instead of assigning each data point to the appropriate subject within each type of measure, the measures were randomly assigned to subjects before the correlations were calculated. This generated an empirical null hypothesis distribution of possible differences for each of the six pairs of correlations. A p-value for each correlation difference was then generated by calculating the proportion of these empirical null distribution values that were as large (or larger) than the difference that was found with the actual (unscrambled) data. Each of these 6 correlation differences were deemed reliable if this proportion was less than .05 (one-sided).

###### Partial correlations

Finally, an alternative to comparing correlations is to perform a multiple regression analysis that includes the nuisance variables within the same model. Therefore, we also fit linear models whose predictors included both the stop-signal P3 *and* each one of the nuisance ERP amplitudes as predictors, with fronto-central P3 to unexpected cues in the CMO task serving as the criterion variable. This produced a partial regression coefficient for the hypothesized correlations between the stop-signal and surprise-related P3 amplitudes, with the influence of the nuisance process (reflected in the control ERP amplitude) factored out.

##### 2.9.2.2. Independent Component Analysis (Approach 2)

Our second, complementary approach to test whether the stop-signal P3 and the fronto-central P3 to unexpected cues reflect overlapping neural processes used ICA.

###### Overview

In all of the analyses above (for both Approach 1 to Hypothesis 2 and for the analyses conducted to test Hypothesis 1), the SST and CMO task data were analyzed separately to avoid any potential bias towards finding a relationship between them. In contrast, for this analysis, the stop-signal and cross-modal oddball data were subjected to the *same* ICA. This allowed us to reanalyze the surprise analyses under Hypothesis 1 with re-constructed data that factored out the signal associated with the stop-signal P3. In this manner, we tested whether the association between the surprise term and fronto-central EEG activity in the cross-modal oddball task relies on the stop-signal IC (suggesting a commonality between processes, 19, 20, 29, 30), or whether processes captured by other ICs explain the surprise-related response in the CMO task (which would suggest that surprise-processing and action-stopping do not involve overlapping processes).

First, we used the SST portion of the data as a functional localizer, extracting one (and only one) independent component (IC) for each subject that best reflected the properties of the fronto-central stop-signal P3. We then generated two different datasets for the CMO task for each subject: one dataset in which the EEG channel data were reconstructed using only the one IC that reflected the stop-signal P3, and one dataset in which the channel data were reconstructed by back-projecting all ICs *except* the stop-signal P3 IC (thereby effectively removing this IC’s contribution from the channel data, similar to ICA-based eye-movement artifact rejection). We then re-ran the single-trial modeling analyses performed under Hypothesis 1, exactly as described above, separately on both datasets.

###### Stop-signal P3 IC selection

Automated selection of the stop-signal IC from the SST portion of the merged data was done using a two-step spatiotemporal selection procedure (31). First, each subject’s component matrix was scanned for components that showed a fronto-centrally distributed positivity on stop-compared to go-trials in the time window 250ms following the respective signal. To this end, the scalp montage was divided into 9 ROIs (an anterior-posterior dimension and a lateral dimension with 3 levels each). Components whose back-projected channel-space topography for that difference wave showed a maximum in the fronto-central ROI (consisting of electrodes FCz, Cz, FC1, FC2, C1, and C2) were selected. From all components that matched this criterion, we then selected the one component whose average time-course across that ROI showed the highest correlation to the original channel-space ERP in the same ROI and time window (i.e., the ERP extracted from a back-projection of all non-artifact components).

###### Stop-signal P3 validation

We reconstructed the channel-space data for both tasks using only the selected component, and tested for the following effects on the SST portion of that dataset to validate that we had successfully selected the stop-signal P3 IC. These tests are direct replications of prior work that established the stop-signal P3 as an index of motor inhibition in the SST (17, 18):

1. The onset of the stop-signal P3 should occur earlier on successful vs. failed stop-trials (as predicted by the race-model of motor inhibition; Logan et al., 1984)
2. The onset of the stop-signal P3 should be positively correlated with SSRT, reflecting its association with the speed of the stopping process.

For these tests, the onset of the stop-signal P3 was quantified as in Wessel & Aron (2015) based on the difference wave between stop- and go-trials (this was done independently for successful and failed stop trials). The time at which the P3 difference wave was largest in the time period 200-400ms following the stop-signal was identified. The analysis worked backwards in time from this maximum difference, with each step backwards occurring only if that step was also significantly greater than 0 (at p < .05). Once a non-significant difference was reached, this determined the time of the P3 onset. The onset times of the successful and failed stop-trials were then compared using a paired-samples t-test (prediction #1 above). Next, the relationship between the successful stop onset and the SSRT across participants was assessed with Pearson’s correlation coefficient (prediction #2 above).

###### Main analysis

We then repeated the model-based single-trial analysis of the CMO task data that was described above for Hypothesis 1, but only on the portion of the EEG data that was explained by the stop-signal P3. In essence, instead of looking at the entire EEG signal, we reconstructed the channel-space signal of the merged EEG data from both tasks by only back-projecting the activity accounted for by the stop-signal P3 IC. We then investigated the task portion of the merged, component-restricted dataset using the same model-fitting procedure as for Hypothesis 1 above. Only the winning model from Hypothesis 1 was fit to the data. If the stop-signal P3 and the surprise-related P3 reflect overlapping neural processes, the model fit should be preserved in that dataset.

Additionally, we also reconstructed a version of the merged dataset that consisted of the back-projection of all original ICs *with the exception of the stop-signal P3 component* (essentially, the inverse of the above dataset). Since participants averaged 17.1 (SEM: .87) components, these data were reconstructed based on the activity of 16.1 independent components. Just as for the single-component dataset that included just the stop-signal P3, we again fit the Bayesian model.

As in Hypothesis 1, this resulted in 640 tests per set (640 for the single-IC dataset and 640 for the other-ICs dataset). As before, the p-values for these tests were corrected across both sets of tests to an alpha-level of .01. This resulted in a corrected alpha-level of p = .00027.

## 2. Results

### 3.1. Behavior

Stop-signal behavior was as expected for a sample of healthy young adults. Mean Go-RT was 520ms (SEM: 15.2), mean failed-stop RT was 444.3ms (SEM:13.2). Mean SSRT was 252.4ms (SEM: 8). Mean error and miss rates were low (1% and 2.6%, respectively). Mean stopping success was 51.4% (SEM: .45, range: 46-59%), demonstrating the effectiveness of the adaptive stop-signal delay algorithm.

In the cross-modal oddball task, correct trial RTs showed the expected pattern as well: There was a main effect of TRIAL TYPE (F(2/108) = 25.3, p = 9.74*10^-10^, partial-eta^2 = .32), a main effect of BLOCK (F(3/162) = 7.64, p = 8.2567*10^-5^, p-eta^2 = .12), and a significant INTERACTION (F(6/324) = 9.78, p = 6.51*10^-10^, p-eta^2 = .15). Individual comparisons revealed that in Block 1, both visual and auditory unexpected-cue RTs were significantly longer compared to standard-cue RTs (t(54) = 9.41, p = 5.48*10^-13^, d = .75 for visual and t(54) = 3.14, p = .0028, d = .29 for auditory, respectively). Furthermore, in Blocks 2 and 4, visual unexpected-cue RTs were also longer compared to standard-cue RTs (t(54) = 4.45, p = 4.3*10^-5^, d = .33 and 3.5, p = .00094, d = .26, respectively). No other comparisons survived corrections for multiple comparisons. Taken together, the data indicate the presence of an initial slowing of reaction times following unexpected cues in both modalities, which wore off over the course of the experiment (Figure 4).

With regards to error rates, there was a significant main effect of TRIAL TYPE (F(2/108) = 3.89, p = .023, p-eta^2 = .067), with no main effect of BLOCK (F(3/162) = .4096, p = .74631, p- eta^2 = .0075), and no INTERACTION (F(6/324) = .7, p = .65, p-eta^2 = .013). The main effect was accounted for by lower error rates on both types of unexpected-cue trials compared to the standard-cue trials, which persisted throughout the task.

**Figure 4.**
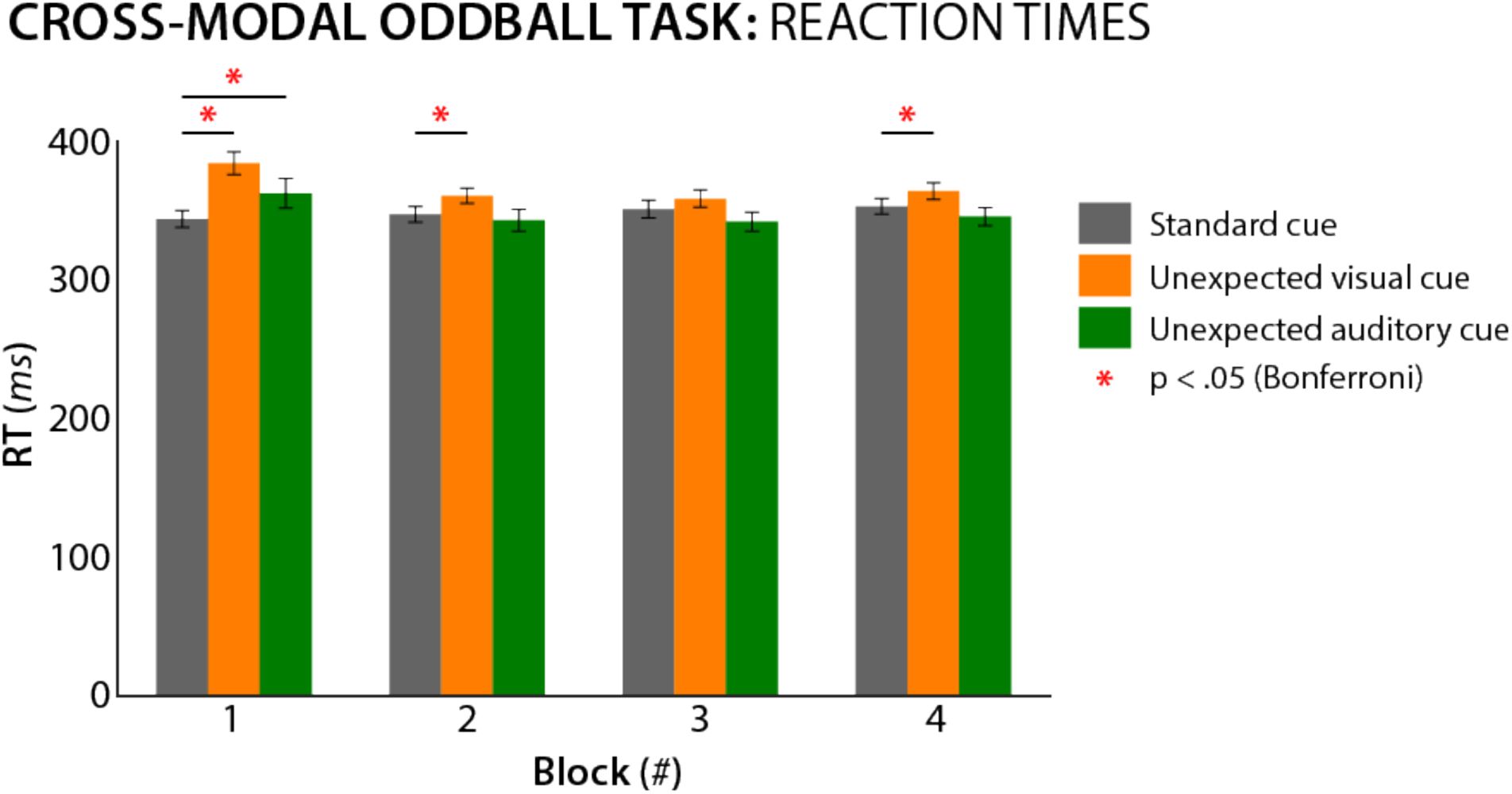
Reaction time data from the cross-modal oddball task. Significant individual comparisons (Bonferroni-corrected) are highlighted in red. For both unexpected auditory and unexpected visual cues, reaction times were slower compared to standard cues in Block 1. This effect wore off over time.

With regards to miss rates, there was no significant main effect or interaction (all p > .14).

### 3.2. Hypothesis 1: Frontal cortex independently tracks surprise depending on sensory domain

#### 3.2.1. Single-trial EEG model fitting

Our single-trial EEG analysis showed that both models significantly fit the data in the time windows centered on 300 and 350ms post-cue (Figure 5). Both model terms show significant positive correlations with fronto-central electrodes, as hypothesized. Positive correlations that exceeded the significance threshold of p = .00044 for Model 1 (separate surprise terms) were found at electrodes Fz, Cz, FCz, FC1, FC2, F1, F2, C1, C2, FC3, and FC4 in the 300ms time window and at electrodes F3, F4, Fz, Cz, FCz, FC1, FC2, F1, F2, C1, C2, FC3, and FC4 in the 350ms time window. For Model 2 (common surprise term), significant positive correlations were found at electrodes Fz, FCz, FC1, FC2, F1, F2, C1, C2, and FC3 in the 300ms time window and at electrodes F4, Fz, Cz, FCz, FC1, FC2, F1, F2, FC3, and FC4 in the 350ms time window.

**Figure 5.**
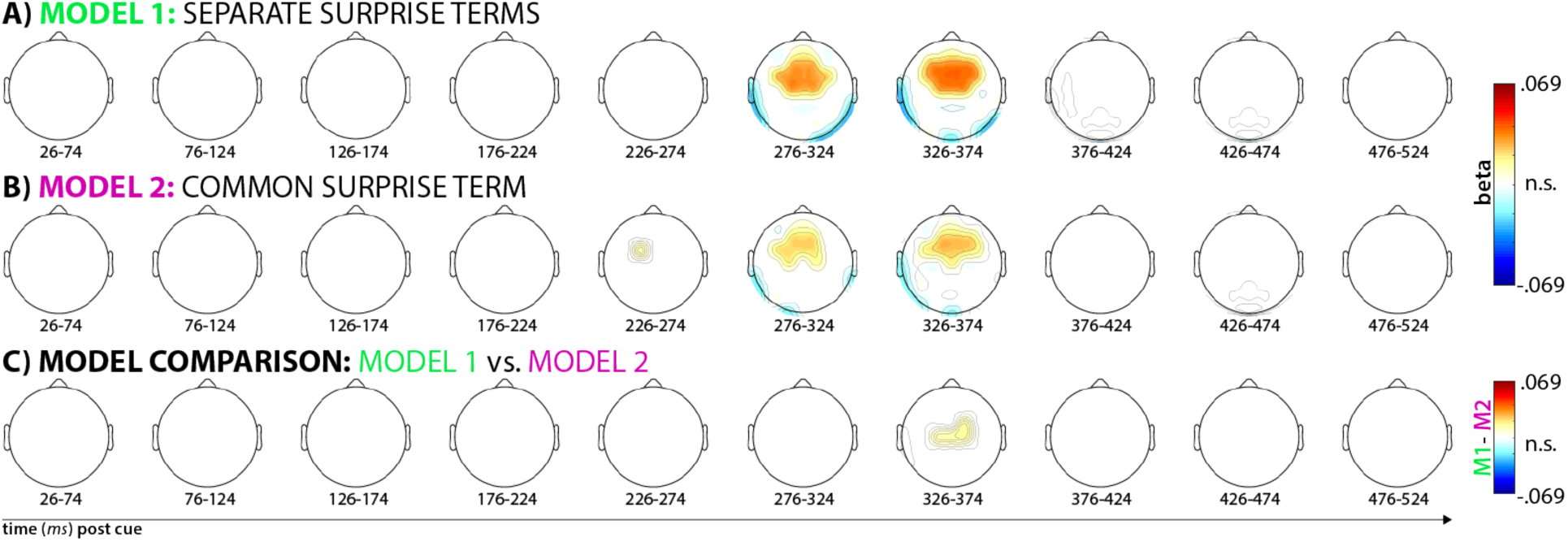
Results from the whole-brain single-trial model fitting analysis described in Figure 3. Each topography depicts the averaged standardized beta coefficient at each channel in the respective time window (x-axis) and model (plots A and B), as well as the M1-M2 model comparison (plot C). White areas denote channels at which the fit within the depicted time-window was non-significant (p < .00044). In A and B, red areas denote significant positive correlations between the respective model and the EEG data, blue areas denote significant negative correlations. In C, red areas denote higher correlations between Model 1 and the data compared to Model 2.

While both models fit the data well at a similar cluster of fronto-central electrodes (which is to be expected, considering that the surprise terms from each model are largely similar), direct model comparisons showed that Model 1 (separate terms) fit the data significantly better than Model 2 (common term). While Model 1 provided numerically better fits at all fronto-central electrodes, the difference was statistically significant at p < .00044 in the 350ms time window at electrodes Cz, FC2, C1, and C2 (Figure 5C).

For illustrative purposes, Figure 6 depicts the trial-averaged time-course of the ERP for all three cue types in a non-windowed fashion at these fronto-central electrodes. As seen in the figure, the time window in which the surprise-model significantly fit the single-trial data (highlighted in beige), both unexpected cues yield a P3 waveform, with the auditory condition producing a noticeably larger deflection.

**Figure 6.**
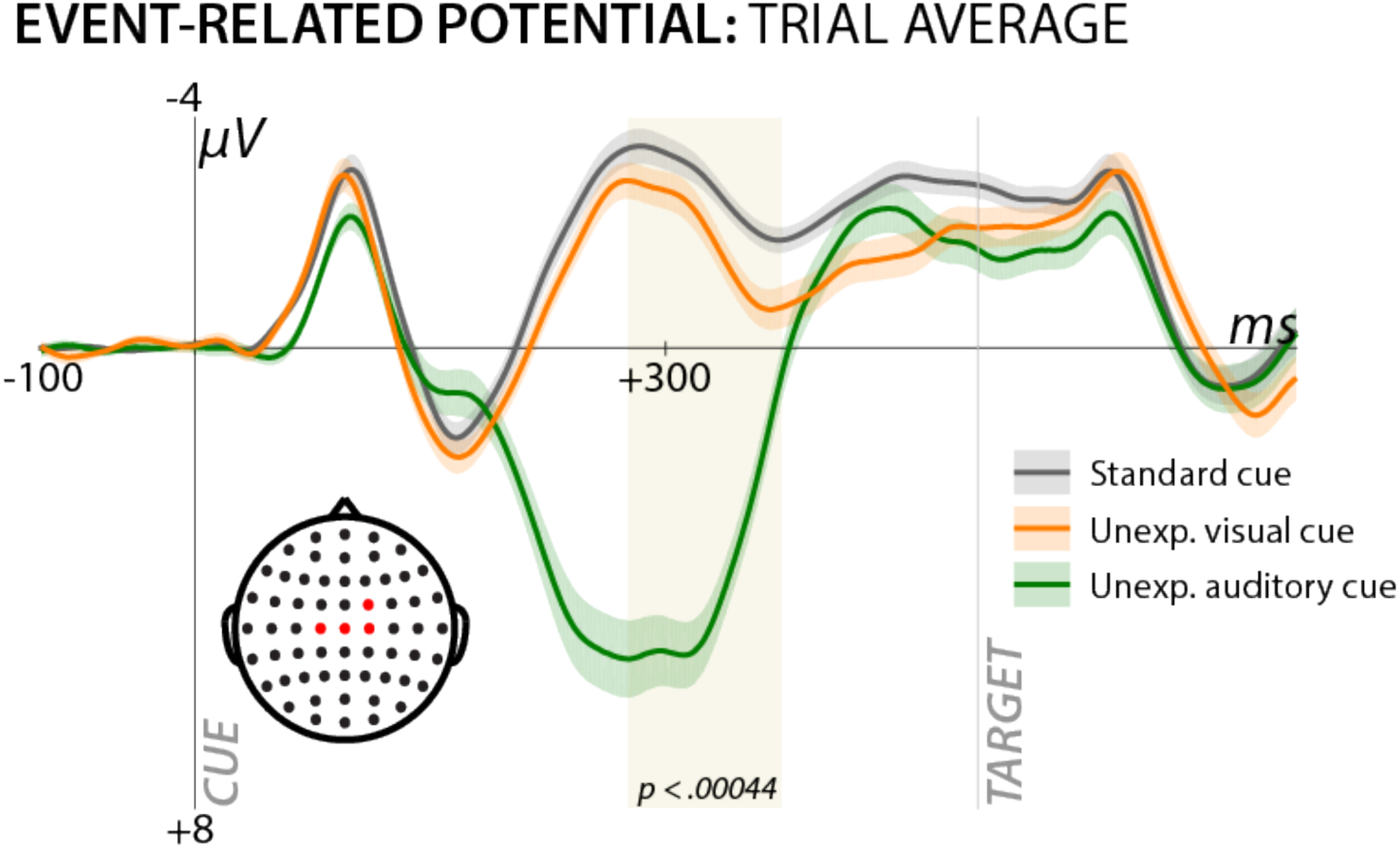
Average channel event-related response to the three different cue types, plotted at the channels in which the winning model (separate surprise terms; Model 1) provided significantly better fit than the losing model (common surprise term). Beige highlighting denotes the time window in which the winning model significantly fit the single-trial EEG response. This trial average illustrates that the time window in which the fit was significant contains the fronto-central P3 ERP to both unexpected auditory and visual cues.

To illustrate that neither sensory domain accounted for the significant model fit on its own, we also plotted the model fits separately for each trial type (rather than using one variable to model both trials types as in the main analysis above). Figure 7 shows the model fits for the separate surprise term (the winning model from the main analysis), split by sensory domain. This revealed that both auditory and visual surprise terms significantly fit the single-trial EEG response to their respective trial type during the same time period and at the same fronto-central scalp sites as the overall fit. This also rules out that the auditory P3, which had a larger amplitude than the visual P3, would solely account for the model fits.

**Figure 7.**
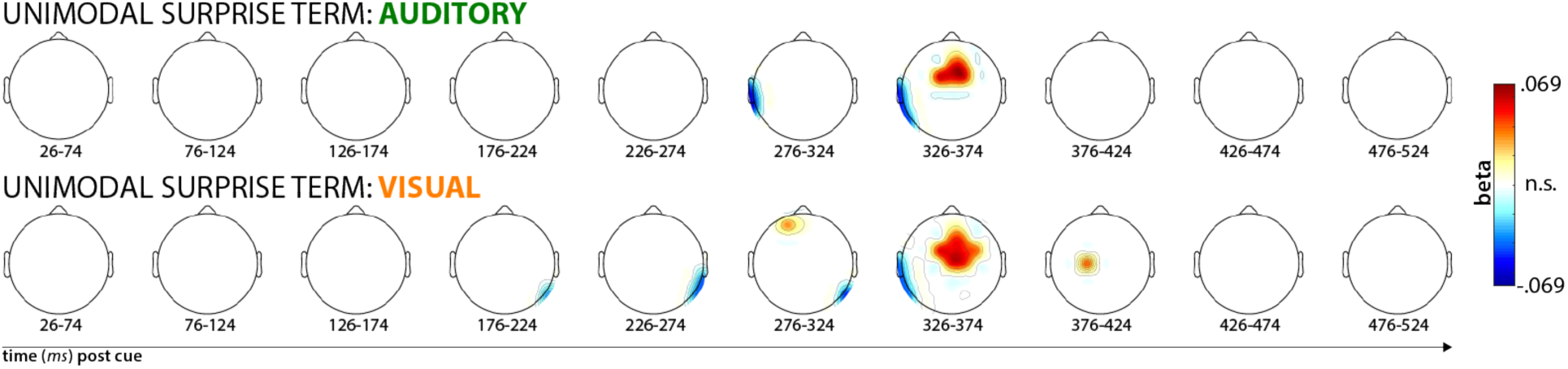
Split model fits of the within-domain surprise values, individually for each sensory domain. Scaling and significance threshold is the same as in Figure 5 (p < .00044). This plot shows that both the auditory and visual unexpected cues contribute to the significant single-trial fit of the separate surprise-terms model in Figure 5A.

#### 3.2.2. Exploratory model-fitting of reaction time latencies

We buttressed our EEG analysis of Hypothesis 1 with an exploratory analysis of the fit between both model terms and each participant’s single-trial reaction times to the target-arrow that followed the cue. Both model terms provided a positive fit with the RT data (i.e., slower RT with surprise), with Model 1 showing a better fit overall, but neither fit was significant at the group level (Model 1: p = .23, Model 2: p = .84). When this analysis was restricted to the first half of the experiment (i.e., the part of the experiment in which the RT effect of the unexpected events had not fully worn off, cf. behavioral results section), both models showed again positive fits between the model terms and RT. For Model 1, the fit was highly significant (t(54) = 3.98, p = .00021), whereas the fit for Model 2 only bordered significance (t(54) = 1.88, p = .066). Just like for the single-trial EEG data, Model 1 (separate terms) fit RT better than Model 2 (combined term); t(54) = 3.71, p = .00049. While this analysis has to be interpreted with caution, given its exploratory nature, it does lend complementary support to the idea that – just like the neural response – the effect of the unexpected cues on behavior is better described by Model 1.

### 3.3. Hypothesis 2: Fronto-central neural activity after surprise indexes inhibitory control

#### 3.3.1. Approach 1: Cross-task ERP amplitude correlations

Figure 8 depicts the correlations between the ERPs across both tasks. In line with our hypothesis that action-stopping and surprise-processing share a fronto-central neural process, there was a significantly positive correlation between the amplitudes of the fronto-central stop-signal P3 in the SST and the fronto-central P3 ERP to unexpected auditory (r = 0.35, p = .0079) and visual (r = 0.35, p = .0087) cues in the CMO.

**Figure 8.**
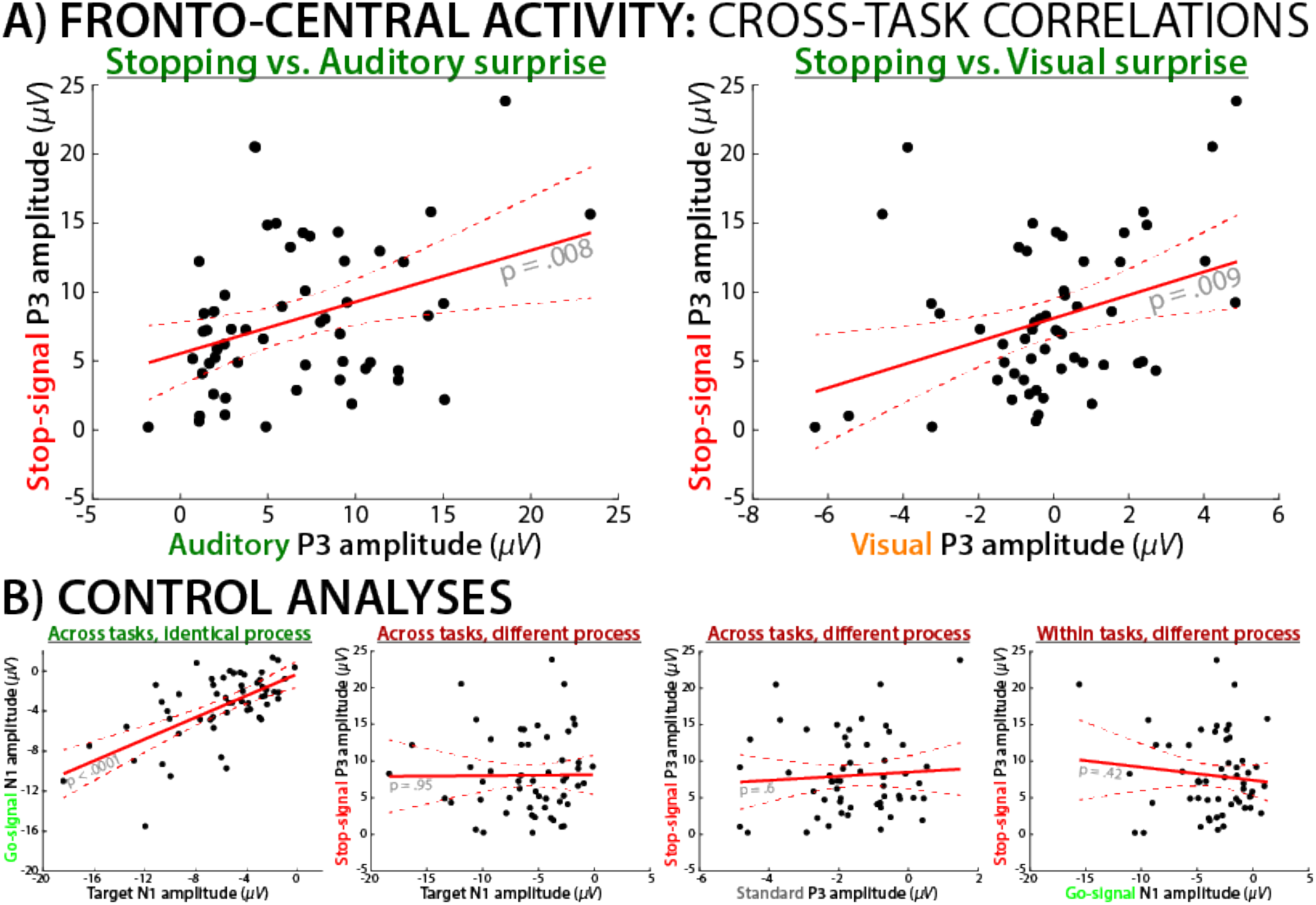
Cross-task correlations between fronto-central activity during surprise processing and action-stopping. A) The amplitude of the stop-signal P3 in the SST was positively correlated with the surprise-related P3 in the CMO task; this was the case for both for auditory (left) and visual (right) unexpected cues. B) Control analyses show that ERP amplitudes of similar processes are indeed positively correlated across tasks (illustrated by the posterior visual N1 to the imperative arrow stimuli in both tasks – i.e., the Go-signal in the SST and the target in the CMO task). Ruling out alternative explanations, there was no reliable correlation for different waveforms from different tasks (middle left; Target visual N1 to stop-signal P3), different waveforms from the same task (right; go-signal visual N1 to stop-signal P3), or the same waveform from different tasks (middle right; standard cue P3 to stop-signal P3). These control analyses demonstrate that there is no general relationship between ERP amplitudes within or across the same task or within the same region of cortex, unless related processes are active.

The control analyses also conformed to our predictions: The posterior visual N1 ERPs to the arrow go-signal in the SST correlated with the visual N1 ERPs to the arrow target in the CMO (r = .55, p = .00001), demonstrating that the same process as occurring in each task can produce a positive ERP correlation. Moreover, there was no significant correlation in any of the control analyses designed to rule out various alternative explanations of the positive correlation between the SST P3 and the CMO P3 (Control analyses 2-4). Specifically, the amplitude of stop-signal P3 was not reliably correlated with the amplitude of the N1 to the Go-signal within the same task (r = -.11, p = .41), demonstrating that individual differences failed to produce a spurious ERP correlation within a task. Similarly, the stop-signal P3 amplitude was not reliably correlated with the visual N1 to the arrow (target) within the cross-modal oddball task (r = .009, p = .95), demonstrating that individual differences failed to produce a spurious ERP correlation across tasks. Finally, stop-signal P3 amplitude was not reliably correlated with the fronto-central P3 amplitude to standard, non-surprising cues in the CMO task (r = .073, p = .6), demonstrating that individual differences failed to produce a spurious ERP correlation for the same ERP component.

In addition to these significance tests on the correlations, our bootstrapping analysis found that the positive correlations between the stop-signal P3 and the fronto-central P3s to unexpected cues in the CMO task were significantly larger than all of the non-significant control analyses. More specifically, the correlation between the stop-signal P3 and the fronto-central P3 to auditory cues was significantly larger than the stop-signal P3 to target-N1 correlation (p = .0136), the stop-signal P3 to go-signal N1 correlation (p = .0482), and the stop-signal P3 to standard-cue P3 correlation (p = .0348). The corresponding p-values for the correlation between the stop-signal P3 and the fronto-central P3 to unexpected visual cues, as compared to the three control correlations were .0144, .0468, and .0373.

Finally, the partial correlation analyses confirmed that the positive correlation between the stop-signal P3 and the fronto-central P3 to unexpected cues in the CMO task could not be accounted for by the amplitude of any of the control ERPs. For unexpected visual cues, the correlation between the fronto-central P3 and the stop-signal P3 was still significant when the model partialed out the Go-signal N1 (partial model fit: t(52) = 2.69, p = .0095), the N1 to the target/arrow in the CMO task (t(52) = 2.7, p = .0094), and the fronto-central P3 to standard cues in the CMO task (t(52) = 3.47, p = .001). The same was true for the correlations between the stop-signal P3 and the fronto-central P3 to unexpected auditory cues (Go-signal N1 partialed out: t(52) = 2.69, p = .0097; N1 to the arrow/target in the CMO task partialed out: t(52) = 2.83, p = .0067; fronto-central P3 to standard cues partialed out: t(52) = 2.7, p = .0094).

#### 3.2.2. Approach 2: ICA

The results from Approach 1 to Hypothesis 2 suggest that action-stopping and surprise-processing involve overlapping neural processes. Providing converging support for this conclusion, we used ICA to investigate whether the trial-by-trial relationship between the Bayesian model surprise terms and the fronto-central activity found in the CMO task was accounted for by the independent component that reflected the stop-signal P3.

We first checked whether the IC that was algorithmically selected to reflect the stop-signal P3 showed the predicted functional properties in the SST (Figure 9). Indeed, the onset of the P3 extracted from that IC occurred significantly earlier on successful stop-trials compared to failed stop-trials (t(54) = 2.4, p = .02, d = .33), and there was a significantly positive correlation between SSRT and P3 onset on successful stop-trials (r = 032, p = .019). Both properties have been previously reported in studies of the SST (e.g., Wessel & Aron, 2015). Hence, we conclude that the selected IC accurately reflected a process that indexes the speed of the motor inhibition process in the SST.

**Figure 9.**
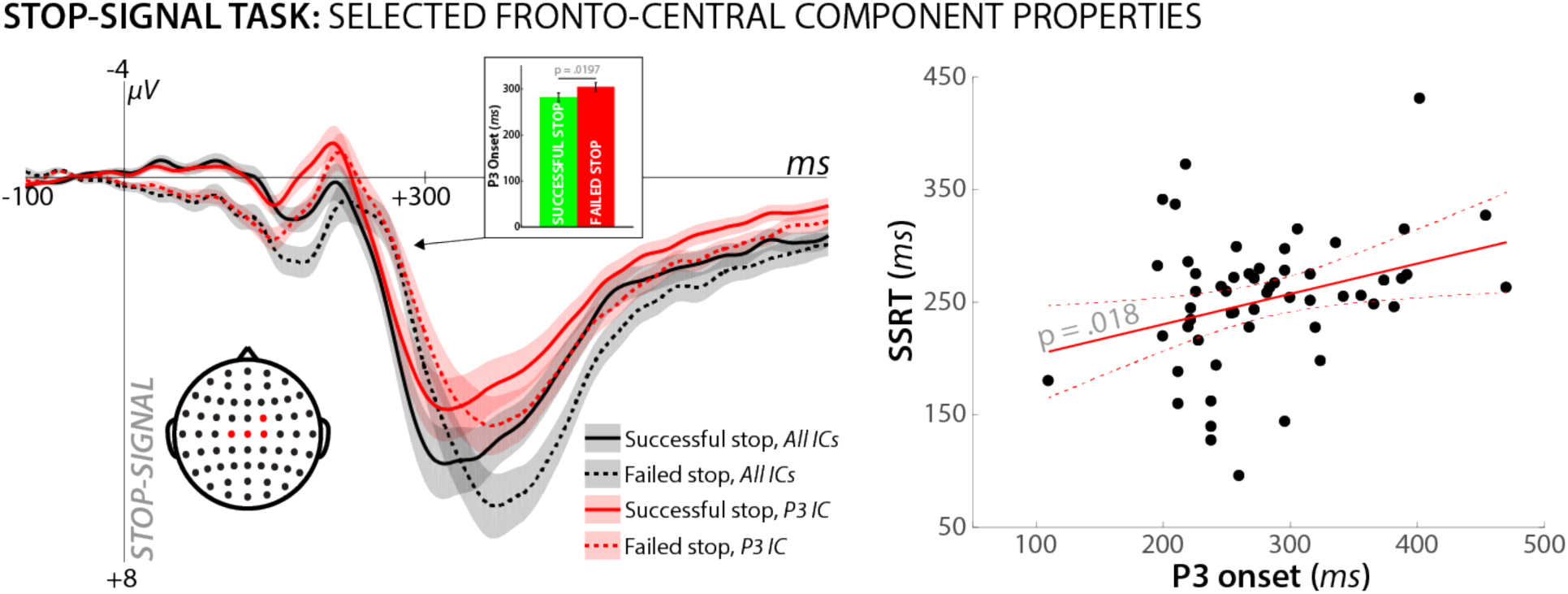
Properties of the independent component selected to reflect the stop-signal P3 in the SST from the merged dataset analysis. The left plot shows that the morphology of the stop-signal P3 based on all non-artifact components (i.e., the standard channel ERP) can be entirely reproduced using just one IC. This shows that the selection algorithm identified the appropriate component, accounting for the activity of the stop-signal P3. Furthermore, this IC shows the classic features demonstrated for the stop-signal P3 in the SST. Namely, the onset of the P3 was earlier on successful vs. failed stop trials (inlay on left plot), and was positively correlated with SSRT across subjects (right plot).

We then repeated our model-fitting analysis (Hypothesis 1) of the CMO task portion of the combined EEG data, when that data was reconstructed using only the selected stop-signal P3 IC for each subject. We found that the winning model from Hypothesis 1 (separate surprise terms) retained its significantly positive fit with fronto-central electrodes (significant positive correlations found in the 300ms time window at electrodes Fz, Cz, FCz, FC1, FC2, CP2, F1, F2, C1, C2, FC4, and C4 and in the 350ms time window at electrodes C4 and F1) when the EEG signal was solely reproduced by back-projecting the stop-signal P3 into channel-space. In other words, the same independent component that indexes successful motor inhibition in the stop-signal task showed the same positive association with the surprise term in the CMO task that was reported for the full channel-space reconstruction (based on all ICs) in Hypothesis 1 (Figure 10A).

**Figure 10.**
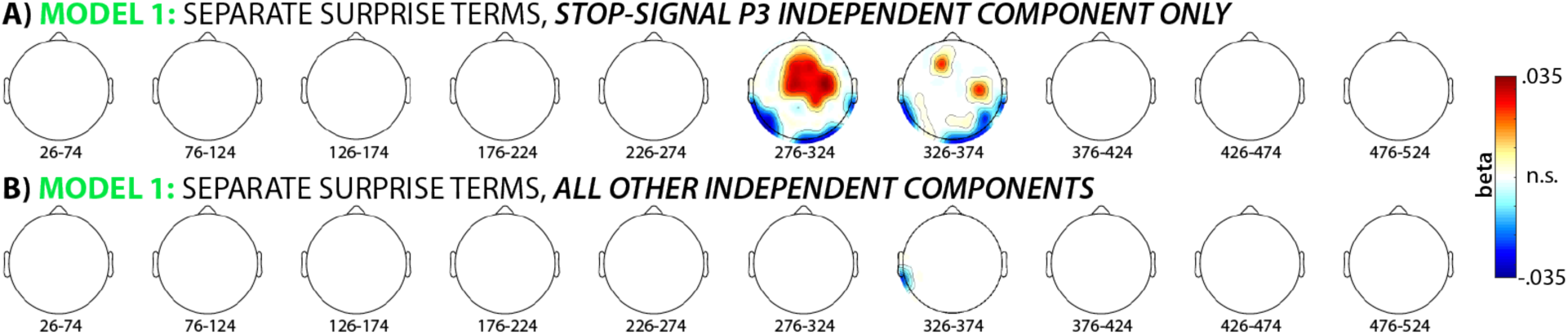
A) A reanalysis of the single-trial model fitting analysis for the winning model (separate surprise terms cf. Figure 5A) using just the one IC that was selected to reflect the stop-signal P3 in the merged dataset. The significant association between fronto-central EEG activity following unexpected cues in the CMO task and the surprise model is retained when the data is reconstructed solely using that one ICA (out of ∼17.1 overall ICs that were extracted per subject on average). B) For comparison, no significant association was found when the data were reconstructed based on the ∼16.1 ICs that did not reflect the stop-signal P3.

In contrast, the remainder of the signal (i.e., the portion of the CMO task EEG data that was reconstructed based on all independent components that were left over after the stop-signal P3 independent component was *removed*) did not show a significant positive association with the surprise term (Figure 10B).

Therefore, we conclude that the same independent component captures stopping-related activity in the SST and surprise-related activity in the CMO task. This confirms the findings of our ERP amplitude analysis in Approach 1 – i.e., that there is overlap between the neural processes following stop-signals (which are not surprising) and surprising cues (which do not instruct the subject to stop).

## 4. Discussion

In the current study, we tested two hypotheses about the nature of surprise processing in human frontal cortex. First, we found that fronto-central event-related activity at roughly 275-375ms following the appearance of unexpected cues tracks surprise for each sensory domain separately. Rather than incorporating surprise into a common cross-modal term, the neural response was better characterized by a model in which surprise was tracked for each domain separately. The time range and topographical extent of this activity overlaps with the well-characterized P3 trial-average ERP, which is in line with classic averaging-based ERP studies of surprise (1, 12, 32). Our single-trial approach was able to disentangle two competing explanations for the common activity found for unexpected events across sensory domains, thereby providing novel insights into how frontal cortex constructs and updates models of the multi-sensory environment.

We then tested whether the modality-independent fronto-central neural activity during surprise indexes a rapid inhibition of ongoing motor activity – i.e., whether the convergence between neural signals following unexpected events, regardless of sensory domain, can be explained by a common control mechanism that is downstream from surprise. This hypothesis is relatively new (15, 33-35), as most previous studies of surprise focused on its cognitive effects (12, 14, 36, 37). The comparatively large sample size of our study allowed us to take the novel approach of correlating electrophysiological signal amplitudes across different tasks, revealing that the P3 amplitude following stop-signals in the stop-signal task reliably correlated with the fronto-central P3 found during multi-modal surprise. Our control analyses indicated that this correlation reflects a common process rather nuisance variables (such as non-specific correlations of ERP amplitudes within or across tasks). Moreover, both ERPs reflected the same component when submitted to a joint independent components analysis.

We conclude that the same process that is underlying the stop-signal P3 is also active during cross-modal surprise. However, what is that process? The most parsimonious explanation is that this signal reflects cognitive control within frontal cortex aimed at inhibiting ongoing motor activity. In the case of stop-signals, this stops the planned motor action, whereas in response to surprise, it produces a ‘pause’, which purchases time for the cognitive system to update the model of the environment without continuing an action that may have been rendered inappropriate by the unexpected change in environmental demand. This pause can also be observed in the reaction time times to the subsequent target. Alternatively, the common process might reflect model updating or surprise (as operationalized in the CMO). However, in the SST, stop-signals are explicitly part of the task (and are introduced during pre-task practice). In other words, participants are *expecting* and planning for stop-signals, and their occurrence should not produce surprise. Indeed, if stop-signals were surprising, one would expect the amplitude of the stop-signal P3 to decrease as the task progressed (i.e., as the priors become stable and the surprise terms become smaller and smaller, which is what occurred for the fronto-central P3 in the CMO task). However, as the auxiliary plot in Figure 11 shows, the amplitude of the stop-signal P3, unlike the P3 to unexpected cues in the CMO task, remained constant throughout the experiment. This is incommensurate with explanations that seek to attribute the cross-task commonalities in neural processing to surprise, infrequency, orienting, or model updating.

**Figure 11.**
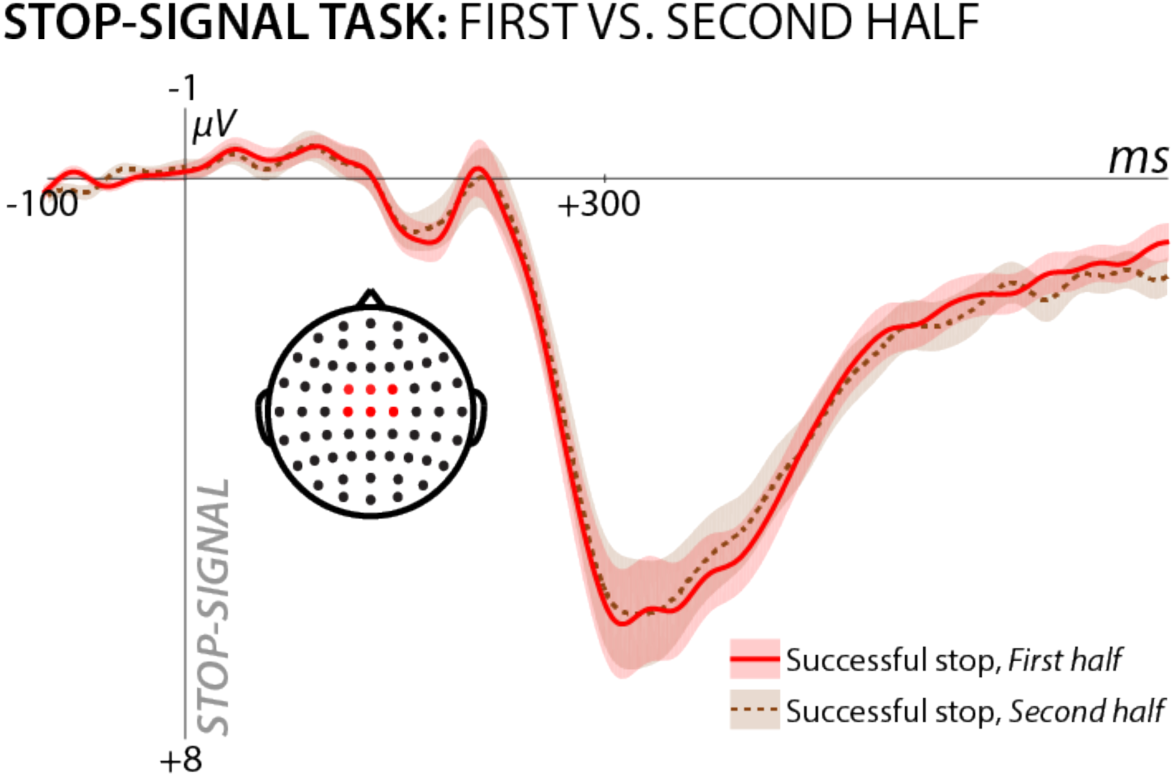
Stop-signal P3 split by phase of the SST experiment. If the process underlying the stop-signal P3 was stop-signal-induced surprise, its amplitude should decrease in the second half of the experiment. Instead, the stop-signal P3 is nearly identical across the two halves of the experiment.

Our preferred interpretation of the common process in terms of motor control is supported by recent studies, which found that unexpected perceptual events lead to a broad, reactive suppression of the motor system, as measured using transcranial magnetic stimulation (35, 38). Additionally, measurements of isometrically exerted force have shown that unexpected events lead to a rapid, reactive reduction of such steadily exerted motor activity (34). Furthermore, unexpected events have been found to interrupt ongoing finger-tapping (39). Finally, studies using optogenetics have shown that when regions of the subcortical network that cause inhibition of motor activity are experimentally inactivated, unexpected events no longer yield interruptive effects on motor behavior (40). All these studies show that surprise, in addition to its prominent cognitive effects, also lead to interruption of ongoing motor activity.

The interpretation that the common process between the stop-signal and CMO tasks is motor control is also supported by some features of our data. Specifically, our behavioral data indicated an incidental slowing of reaction times to the target in the CMO task when that target was preceded by unexpected cues, which is in line with prior behavioral studies (41-43). Our exploratory analysis showed that during the task period in which this RT effect was present, the surprise model (specifically, the separate-term model that also provided the best fit to the neural data) was positively related to the RT data: trials with more surprising cues, according to the Bayesian model, yielded longer reaction times to the subsequent target. We propose that this extra time reflects a momentary suppression of the motor system produced by the unexpected event. Supporting this claim that this ‘pause’ is an adaptive process, accuracy was also increased following unexpected cues (i.e., a speed-accuracy tradeoff was enacted after unexpected cues, which may be enabled by the transient pause in the motor system that we purport to be reflected in the fronto-central P3). In that vein, one notable observation is that while the surprise term fit the neural data for both domains to similar degrees (Figure 7), the trial-average response to unexpected auditory cues in our current study appeared to be larger in amplitude compared to unexpected visual cues (Figure 6). Interestingly, the reverse was the case in the reaction time pattern, where visual unexpected cues seemed to have larger effects (Figure 4). While we are hesitant to make strong conclusions based on the trial-average data, it is notable that the timing of the P3 to the different stimuli also differs in latency, which likely reflects the fact that early auditory processing is faster than visual processing (44). Since the increase in trial-averaged P3 response to unexpected visual cues extends to a time period much closer to target presentation (compared to the P3 to auditory unexpected cues, cf. Figure 6), it is tempting to assume that this may explain the difference in RT effects. However, further studies are necessary to explicitly test this hypothesis.

There is some debate in the literature about the interpretation of the surprise term used in our model comparison analysis. We followed the nomenclature of Itti and Baldi (2010), who termed the calculation of the Kullback-Leibler divergence of the posterior and prior probability distributions (Equation 1) ‘Bayesian surprise’. However, other authors have interpreted this term as ‘model updating’, rather than surprise (45). Instead of KL divergence, they favor Shannon-based information theoretical quantifications of surprise (i.e., surprise is quantified as the inverse of the log-scaled prior expectation of a given stimulus, 46). In past EEG studies, such Shannon-based surprise has been related to the amplitude centro-parietal P3 ERP (47, 48), rather than the fronto-central P3. This is in line with BOLD activation of parietal cortex, which tracks such Shannon-surprise in fMRI (45). Conversely, trial-by-trial indices of Bayesian surprise are associated with the fronto-central P3 (48), which is in line with the current study, as well as with fMRI work showing that BOLD activity in medial frontal cortex tracks Bayesian surprise (45). Collectively, these results underscore that Shannon-surprise and Bayesian surprise are not only different computational terms but that they may be related to different neural signals.

However, in terms of the theoretical distinction between Bayesian surprise and Shannon surprise, it is important to note that both concepts are closely related in most circumstances – i.e., whenever there is surprise, it will lead to the updating of internal models of the environment. This is also reflected in a high correlation between Shannon- and Bayesian surprise that is present in most experimental circumstances (including the current one). Under some circumstances, it is possible to untangle surprise and model updating by introducing different degrees of volatility into the environment (49) or by explicitly instructing participants that certain surprising cues should not be used to update the internal model of the task (45). However, in studies like the current one, the two terms are largely identical, with the exception being trials in which in an unexpected cue follows a prolonged sequence of expected cues. (Such trials introduce non-monotonous upticks in the Shannon surprise term, whereas the Bayesian surprise / model updating term is always monotonically decreasing). Perhaps most relevant is the question which term better reflects the commonplace meaning of ‘surprise’ in the everyday world, outside of the laboratory, and which term better reflects the participants’ approach to the experiment. If subjects place strong emphasis on the recent trial sequence and dynamically adapt to the changing local probabilities of unexpected cues, then the Shannon term may provide a better characterization of surprise. This would be the case if participants assume that the current environment constantly changes (i.e., high volatility). However, if subjects approach the experimental task as a specific, unchanging environment that they need to adapt to by learning the base rates of occurrence, then the Bayesian surprise term may provide a better characterization of surprise. In the current study we assumed that the latter is the case (indeed, the experimental design involved a stable procedure for each task), and as such, ‘surprise’ and ‘model updating’ are essentially synonymous in our study.

Taken together, our study suggests that when an environmental model is updated because of an unexpected cue, this leads to surprise, which is accompanied by inhibitory control of the motor system. From a real-world perspective, it makes sense for the cognitive apparatus to operate this way. Because we interact with the environment by executing motor commands, it is important that we interrupt ongoing motor behavior while the model of the environment is updated; ongoing actions need to be re-evaluated in light of changing environmental contingencies. We hypothesize that motor inhibition prevents the execution of actions that were appropriate under the old, now outdated model, and may also free up resources to rapidly initiate appropriate new actions. This interpretation of the medial frontal cortex is in line with prior findings regarding its role in the control of behavior (2, 50, 51). Here, we propose a specific neural mechanism by which such control of behavior is achieved during surprise.

In conclusion, we found that surprise-based model updating in frontal cortex occurs separately for each sensory domain, but shares a supra-model control mechanism that likely involves the inhibitory control of behavior. These results suggest a specific control mechanism that is rapidly deployed when the model of the environment unexpectedly changes.

## Acknowledgements

The authors thank the following individuals for help with data collection: Hailey Billings, Nathan Chalkley, Kylie Dolan, Isabella Dutra, Tobin Dykstra, Jenna Kelly, Cailey Parker, Julian Scheffer, and Darcy Waller. This research was funded by the following grants: NIH R01 NS102201 and NSF CAREER 1752355 to JRW; NIH RF1 MH114277 to DEH.

## Footnotes

1 This formulation of Model 1 assumes statistical independence in calculating these probabilities. However, the two kinds of unexpected cues were not statistically independent in the experimental design, as no trial included unexpected cues for both sensory domains. To address this, we investigated an alternative formulation of Model 1 that respected this mutual exclusivity inherent in the experimental design. For example, upon realizing that the current trial contained an expected visual cue, this increases the prior probability for an unexpected auditory cue. It is not clear whether subjects could have reasonably learned this mutual exclusivity. Regardless of the formulation of Model 1, the first unexpected event for either modality is maximally surprising (as a result, this alternative formulation of Model 1 was nearly identical to the reported version, which assumed statistical independence).

